# Podoplanin expression identifies human airway basal cells with higher progenitor cell potential

**DOI:** 10.64898/2025.12.17.694989

**Authors:** Sarah E. Clarke, Charlotte C. Percival, Adam Pennycuick, Zoe E. Whiteman, Yuki Ishii, Sandra Gómez-López, Maral J. Rouhani, Ahmed Alhendi, Jessica C. Orr, Hugh Selway-Clarke, Marie-Belle El-Mdawar, Zoe C. Hagel, Kate E. J. Otter, Chuen R. Khaw, Helen Hall, Amyn Bhamani, Robert E. Hynds, Kate H.C. Gowers, Sam M. Janes

**Author notes:** **Correspondence:** Sam M. Janes.

## Abstract

Basal cells are key to maintaining and repairing a functioning airway epithelium. Understanding how basal cells maintain normal airways provides a foundation for interpreting their dysfunction in disease states and for the development of novel therapies. The airway epithelium exists in a dynamic state in which basal stem cells replace lost luminal mucosecretory and multiciliated cell types via an intermediate ‘suprabasal’ cell state. The ability to isolate basal cells with high progenitor cell potential would be beneficial in regenerative medicine applications, but the molecular identity of this population is unclear. Here, we evaluate candidate surface markers to isolate human basal cells. As an individual marker, we found that podoplanin (PDPN) had a favorable sensitivity and specificity compared with integrin alpha 6 (ITGA6) or nerve growth factor receptor (NGFR). We found that KRT5-expressing basal cells could be subdivided into those with high or low PDPN expression; KRT5-negative cells did not express PDPN. *In vitro*, PDPN-high basal cells had higher colony-forming capacity, increased population doubling potential and formed larger colonies than PDPN-low basal cells. PDPN-high basal cells expressed higher levels of *TP63*, as well as other genes expressed by ‘quiescent’ or ‘resting’ basal cells identified in single cell RNA sequencing studies. PDPN-low basal cells expressed genes associated with a ‘differentiating’ basal cell state, including *KRT4*, *NOTCH3* and serpin B family genes. Our results demonstrate that PDPN expression can identify basal cells with high progenitor cell potential, enabling high efficiency sorting of airway stem cells.

## Introduction

The human airway epithelium is a pseudostratified tissue that extends from the nasal cavity to the small airways through multiple generations of conducting bronchi [1]. Its luminal surface comprises ciliated and mucosecretory cells, which cooperate to maintain mucus production and microbial clearance. Basal cells, which reside along the basement membrane, play a key role as airway epithelial stem/progenitor cells, supporting tissue homeostasis and regeneration after injury [2,3].

Basal cells can be identified by the expression of various intracellular marker proteins, including the intermediate filament protein keratin 5 (KRT5) and the transcription factor tumor protein p63 (TP63) [4,5]. Lineage-tracing experiments in mice suggest that there are a continuum of differentiation states among Krt5-expressing basal cells. Less differentiated cells are Krt5+/Krt8-, while Krt5+/Krt8+ ‘suprabasal’ or ‘intermediate’ cells are more committed to differentiation [6,7]. Similarly, single cell RNA sequencing analyses of human airway epithelium identify basal cells in a range of differentiation states. Although specific cluster annotation and marker genes vary between studies, data point to the coexistence of populations of quiescent basal cells, differentiating basal cells and proliferating basal cells. The quiescent, or ‘resting’, basal cells are identified by high *TP63* expression, while the differentiating suprabasal cell populations express members of the serpin B family, along with *NOTCH3, KRT4* and *KRT13* [8–10]. ‘Cycling’ or ‘proliferating’ basal cells are identified by high expression of *MKI67* and other cell cycle-associated genes.

Airway epithelial cell culture studies support the notion that KRT5+ cells are heterogenous in their stem cell potential. Mouse studies confirm that a subset of Krt5-expressing basal cells have different *in vitro* colony forming potential [4], but, as KRT5 is intracellular, live human KRT5 cells cannot be sorted for study and there is little consensus regarding optimal basal cell isolation strategies in the literature. Various surface proteins have been used to identify basal cells, including integrin alpha 6 (ITGA6), nerve growth factor receptor (NGFR) and podoplanin (PDPN) [15,17–23]. ITGA6 is enriched in basal cells [22] and is a component of hemidesmosomes, linking the intermediate cytoskeleton to the extracellular matrix. NGFR, a member of the tumor necrosis factor receptor superfamily that binds to neurotrophins, is also more highly expressed in basal cells than their differentiated progeny [20]. PDPN (also known as T1*α*) is a mucin-type transmembrane glycoprotein that is expressed in a variety of tissue and cell types, including lymphatic endothelium, alveolar type 1 cells and mesothelial cells [25]. PDPN was used in combination with CD166 (ALCAM) and CD49f (ITGA6) to isolate different groups of airway epithelial cells, with CD49f^hi^T1α^+^CD166^mid^ cells expressing relatively higher levels of KRT5 and TP63 [15].

In this study, we aimed to evaluate basal cell surface markers for their ability to identify basal cells. We then sought to understand how we may isolate and study subsets of basal cells to investigate if basal cell heterogeneity, as purported by single cell transcriptomics, extended to discernable functional differences *in vitro*. Studying human airway basal cell transcriptomic and functional differences *in vitro* will help further our understanding of airway stem cell dynamics. The potential to isolate basal cells with high progenitor cell potential could be particularly useful in the context of stem cell therapies and airway epithelial regeneration and repair.

## Results

### Podoplanin is a sensitive and specific extracellular marker of human airway basal cells

We compared the sensitivity and specificity of ITGA6, NGFR and PDPN as extracellular surrogate markers for KRT5 in freshly dissociated second and third generation human airways using flow cytometry. EPCAM was used as a pan-epithelial cell marker, while endothelial (CD31+) and immune cells (CD45+) were excluded from analysis (henceforth EPCAM+ cells). Proximal airway cells were dissociated into a single cell suspension using dispase and trypsin and the expression of ITGA6, PDPN and NGFR was evaluated among KRT5+ and KRT5-EPCAM+ cells (sorting strategy demonstrated in Supplementary Figure 1). ITGA6 and PDPN marked 95.6% and 85.6% of KRT5+ cells, respectively (Figure 1A), while NGFR marked just 11.2%. Concerned that the dispase/trypsin digestion might cleave NGFR, we repeated the analysis using liberase only digestion and found that NGFR marked 55.5% of KRT5+ cells (Figure 1A). Of the KRT5-cells, ITGA6 marked 39.2% of cells, PDPN 15.2% and NGFR (using liberase digestion) 10.7% (Figure 1B). Since PDPN identified almost as many KRT5+ cells as ITGA6 but fewer KRT5-cells, we judged that PDPN expression provides the best balance between sensitivity and specificity of these three putative basal cell markers.

**Figure 1:**
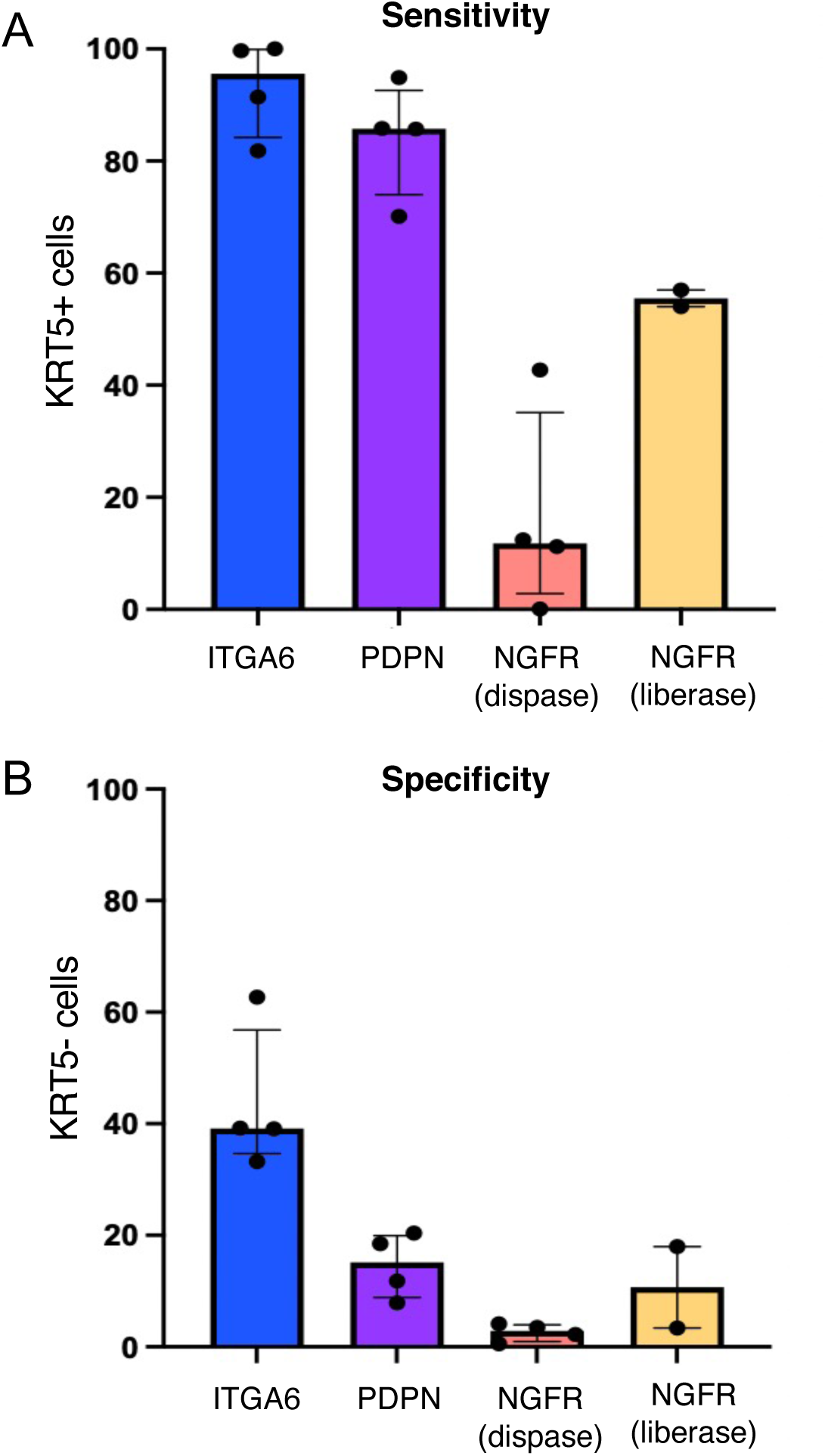
Podoplanin is a sensitive and specific extracellular marker of basal cells in the proximal human airway. A) The proportion of CD45/31-, EPCAM+, KRT5+ cells that express the candidate extracellular markers, ITGA6, PDPN and NGFR, as determined by flow cytometry. B) The proportion of CD45/31-, EPCAM+, KRT5-cells that express the candidate extracellular markers, ITGA6, PDPN and NGFR, as determined by flow cytometry. For both A and B, the bar represents the median and error bar shows the interquartile range. The points represent biological replicates.

### scRNAseq data highlights the variability in expression of basal markers in basal cell subsets

To investigate the expression patterns of *PDPN*, *TP63*, and *KRT5* in basal, differentiating basal cells and terminally differentiated cells of the human trachea, we used single-cell RNA sequencing data from a recent study [16]. Cells were grouped into broad cell-type annotations; basal cells (*n* = 22,580), differentiating basal cells, comprising KRT13/KRT4 and suprabasal populations, (*n* = 13,344), and terminally differentiated cells, which included all secretory cells (*n* = 5,806; Figure 2A). Visualization of the expression profiles of *PDPN*, *TP63*, and *KRT5* revealed that *PDPN* and *TP63* were most highly expressed in basal cells (Figure 2B; Supplementary Figure 2). While *PDPN* and *TP63* expression were strongly associated with basal cells, *KRT5* expression was found to be expressed across basal cells and differentiating basal cells (Figure 2B & 2D). The lowest expression of *PDPN*, *TP63*, and *KRT5* was amongst the terminally differentiated cells (Figure 2B & 2D). This observation suggests that *KRT5* expression is not as exclusive to undifferentiated basal cells as *PDPN*, highlighting the differential specificity of these markers in the airway epithelium.

**Figure 2:**
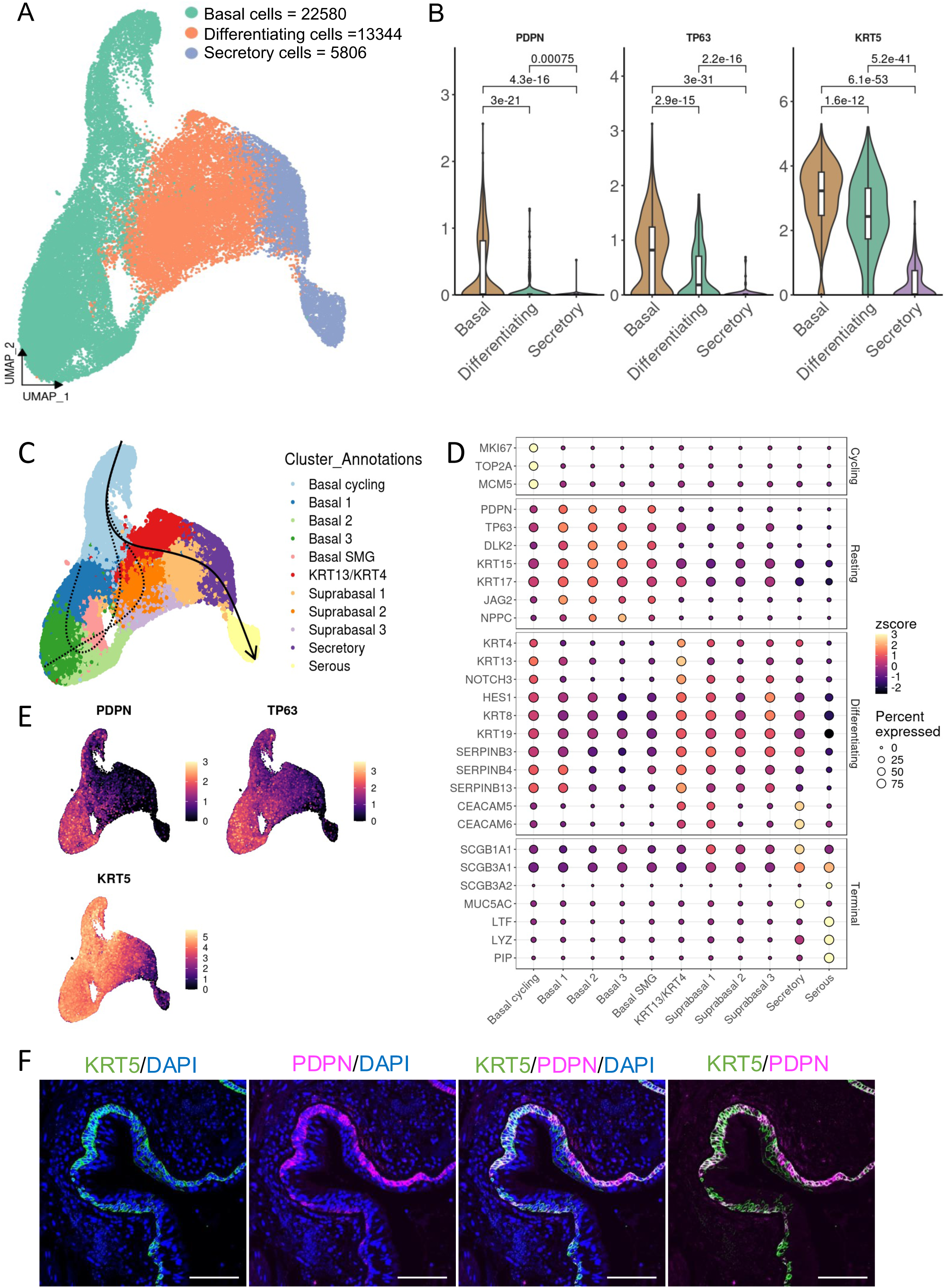
Podoplanin mRNA expression marks resting basal cells in the human trachea. A) UMAP visualization of epithelial cells in the human trachea (*n* = 41,730). Clusters are colored according to broad cell-type annotation. B) Violin plots showing expression levels of *PDPN*, *TP63*, and *KRT5* across broad epithelial cell types. P-values are adjusted using the Benjamini–Hochberg method (Wilcoxon test). C) Slingshot trajectory inference, with cycling basal cells set as the starting point for pseudotime. Dashed lines indicate trajectories towards basal and suprabasal subpopulations, while the solid line represents the lineage from cycling basal to secretory cells. D) Grouped dot plots illustrating the expression of cycling, resting, differentiating, and terminal markers across epithelial cell types in the human trachea. E) UMAP visualizations of gene expression for *PDPN*, *TP63*, and *KRT5*. F) Immunofluorescence staining of proximal human airway epithelium demonstrates co-expression of KRT5 with PDPN along the basement membrane in three biological replicates at 20x magnification. Scale bar = 100 μm.

To further explore the differentiation trajectory, we employed Slingshot for trajectory analysis (Figure 2C). This analysis revealed that *PDPN* expression is highest amongst resting basal cells. In addition, we identified 10 novel genes (*IFFO2, CLSTN1, PGD, AURKAIP1, SDHB, DHRS3, TCEA3, RPL11, SRSF10, and ZNF593*) that displayed significant changes in expression over pseudotime (data not shown). These genes have not been previously reported in the context of airway epithelial differentiation and may represent novel markers of intermediate cell states between basal cell and secretory lineages.

Immunofluorescence staining for KRT5 and PDPN expression in proximal human airway specimens from three biological replicates showed that PDPN is co-expressed with KRT5 in cells along the basement membrane (Figure 2F; Supplementary Figure 3).

### KRT5+ basal cells can be subdivided into two populations on the basis of podoplanin expression

Among freshly flow sorted EPCAM+ cells, the PDPN+ population comprised two subpopulations of cells: those that expressed a low level of PDPN, termed PDPN-low cells, and those that expressed higher levels of PDPN, PDPN-high cells (Figure 3A). There was also an EPCAM+, ‘PDPN-negative’ population (Figure 3A). The proportions of these populations amongst EPCAM+ cells were consistent over multiple donors: the PDPN-negative population represented 67.6% of EPCAM+ cells, the PDPN-low 6.8% of EPCAM+ cells and the PDPN-high 9.4% of EPCAM+ cells (Figure 3B). Taken together, the proportion of PDPN-positive (PDPN-low and PDPN-high) cells is consistent with literature suggesting that basal cells comprise around 30% of the proximal airway epithelium [26, 27].

**Figure 3:**
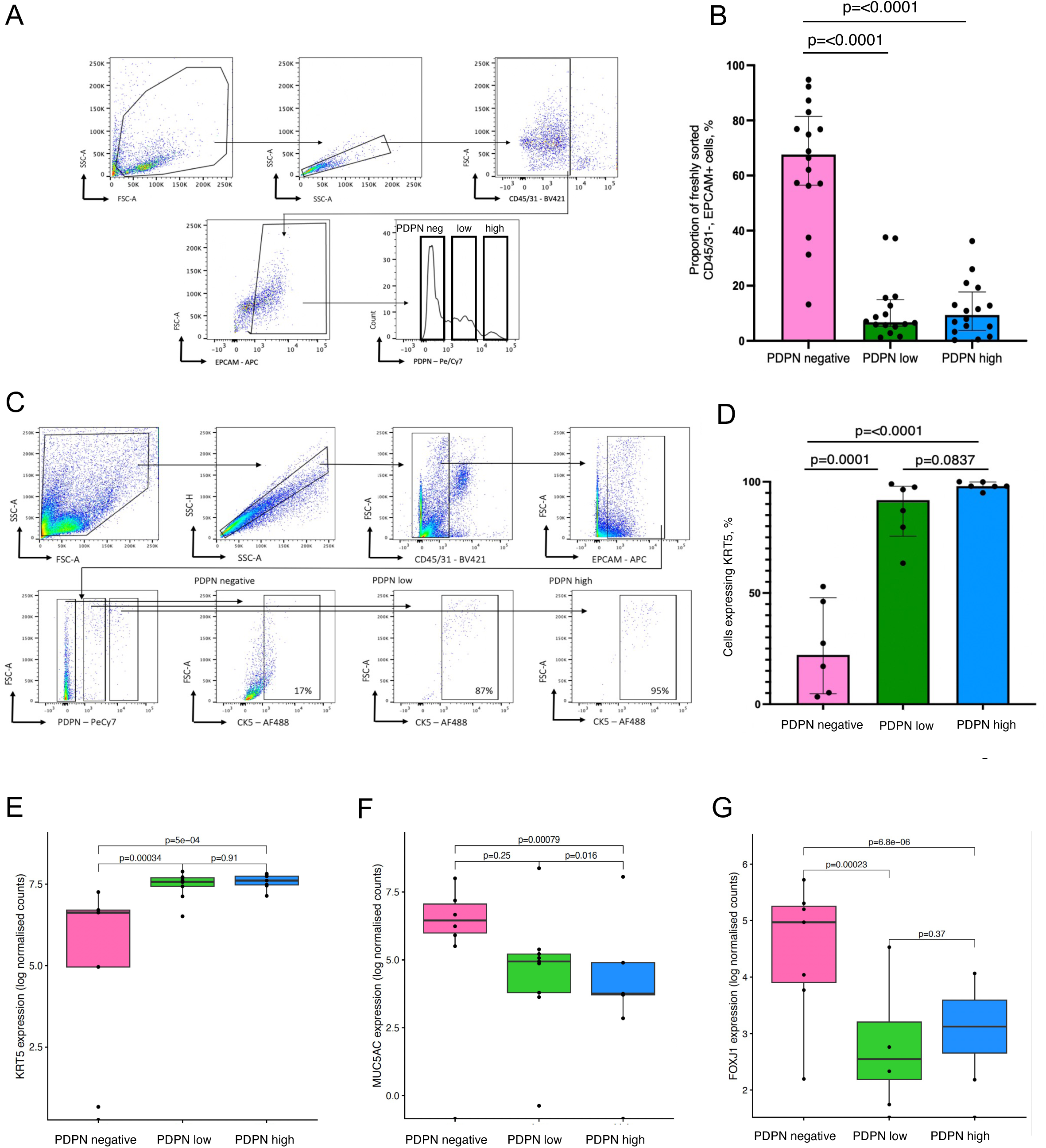
Basal cells can be subdivided into two populations on the basis of podoplanin expression. A) Fresh human proximal airway samples were digested and analyzed using flow cytometry. Representative flow cytometric analysis from one proximal airway specimen demonstrating the pattern of PDPN expression for CD45/31-, EPCAM+ cells. Three populations of cells are identified based on level of PDPN expression: PDPN-negative, PDPN-low and PDPN-high cells. B) Bar chart showing the median proportion of PDPN-negative, -low and –high cells amongst freshly digested and flow-sorted CD45/31-, EPCAM+ airway samples. The error bars indicate the interquartile range. Each point represents an individual biological replicate. One-way ANOVA test comparing all three groups showed a significant p-value (p=<0.0001) so individual unpaired t-tests were carried out for which the p-values are shown. C) Representative flow cytometric analysis of KRT5 expression in CD45/31-, EPCAM+ cells gated based on level of PDPN expression. D) Bar chart shows the median percentage of KRT5+ cells among CD45/31-, EPCAM+, PDPN-negative, -low and -high populations from the flow cytometric analysis of proximal airway from 6 biological replicates. The error bars indicate the interquartile range. A one-way ANOVA compares all three groups (p = <0,0001). Individual unpaired t-tests between groups were carried out for which p values are shown. Levels of *KRT5* (E), *MUC5AC* (F), *FOXJ1* (G) expression in bulk RNA sequencing analysis of flow-sorted CD45/31-, EPCAM+, PDPN-negative, -low and -high cells from freshly digested proximal human airway. The box plot represents the median and interquartile range. The whiskers extend to the smallest and largest values no further than 1.5 IQR from the hinge. Each point represents individual biological replicates. The results of Wilcoxon ranked tests are shown.

We next assessed the levels of KRT5 expression in PDPN-negative, PDPN-low and PDPN-high cells in freshly isolated cells (Figure 3C). By flow cytometry, 98% of PDPN-high cells and 91.8% of PDPN-low cells were KRT5+ (Figure 3D), while only 22.2% of PDPN-negative cells were KRT5+ (Figure 3D). Bulk RNA sequencing of fresh, flow-sorted PDPN-negative, PDPN-low and PDPN-high cell populations further demonstrated *KRT5*+ expression was significantly higher in the PDPN-low and PDPN-high populations compared to the PDPN-negative cell population (Figure 3E). Expression of *MUC5AC* and *FOXJ1*, markers of differentiated mucosecretory and ciliated cells, respectively, was higher in the PDPN-negative cell population than either the PDPN-low or PDPN-high cell populations (Figure 3F-G). Both flow cytometry and RNA sequencing data provide support for the notion that PDPN-low and PDPN-high cells are basal cells, while the PDPN-negative population predominantly contains differentiated airway epithelial cell types.

### PDPN-high cells express genes associated with quiescent basal cells

To identify potential differences in the transcriptomes of PDPN-low and PDPN-high cells, we further explored bulk RNA sequencing data from freshly sorted epithelial cells. Principal component analysis (PCA) showed that, at a group level, the PDPN-low and PDPN-high cell populations were the most closely related, and could be distinguished from the PDPN-negative population (Figure 4A). Donor-donor variability was greatest amongst the PDPN-negative cells (Figure 4A). Hierarchical clustering also pointed to similarities in gene expression between PDPN-low and PDPN-high cell populations, which tended to cluster together (Figure 4B). From the unbiased variance analysis depicted as a heatmap, the genes differentially upregulated in the PDPN-negative group included the mucins *MUC5B*, *MUC5AC* and *MUC7*, as well as the surfactants, *SFTPC*, *SFTPB* and *SFTPA1* (Figure 4B). Additional genes associated with differentiated cell types that were upregulated in the PDPN-negative population included ciliary structural proteins, such as *SNTN* and *DNAH5*. These findings are consistent with the PDPN-negative population predominantly containing differentiated mucosecretory and ciliated cells.

**Figure 4:**
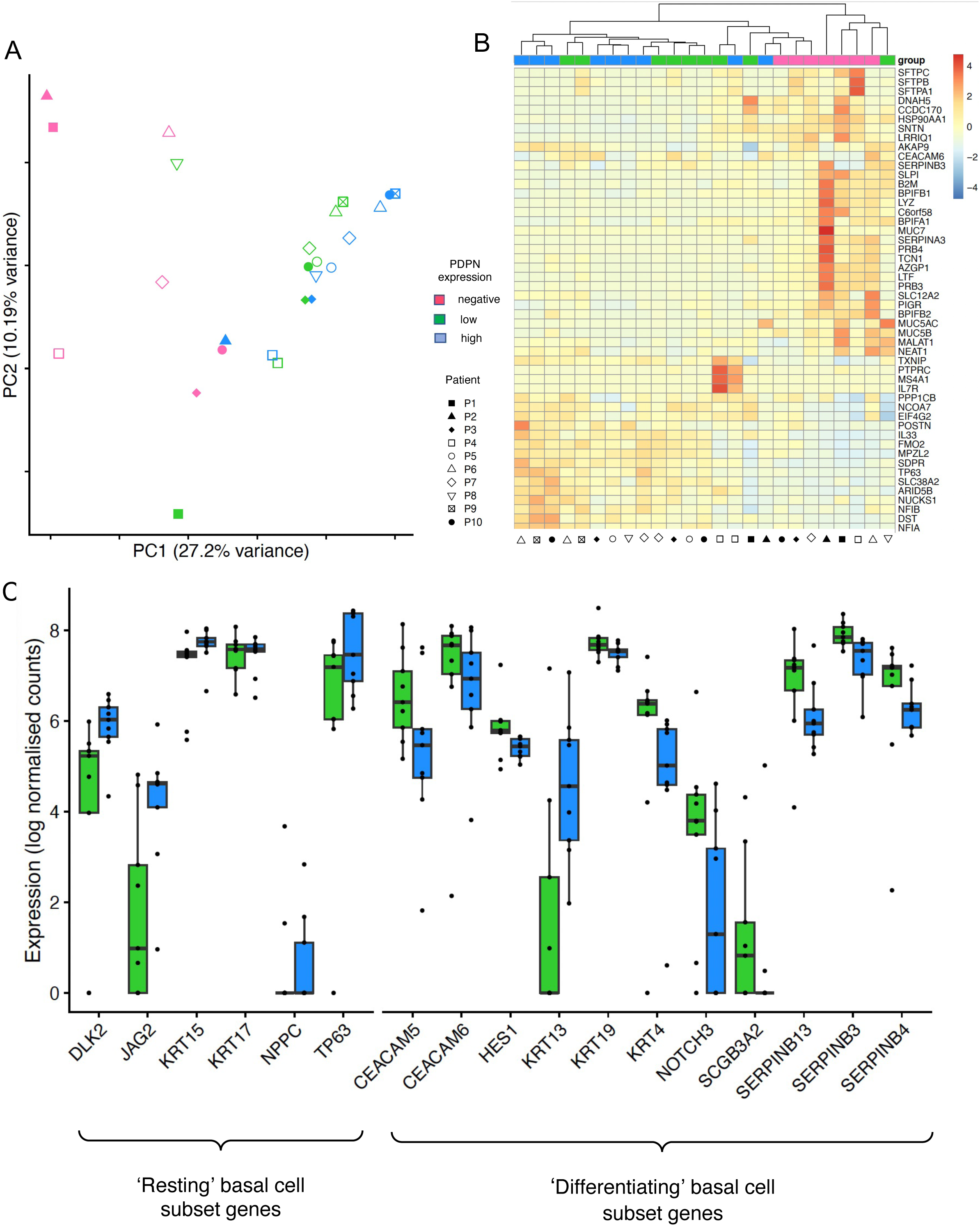
Transcriptomic data shows ‘PDPN-high’ basal cells express genes associated with a ‘resting’ basal cell state and ‘PDPN-low’ cells express genes associated with a ‘differentiating’ basal cell state. A) Principal component analysis showing principal component (PC) 1 in relation to PC2 of bulk RNA sequencing data from freshly flow sorted CD45/31-, EPCAM+, PDPN-negative, -low and -high cell populations. Individual points represent cell populations from each donor and are colored according to level of PDPN expression. B). Unbiased variance analysis of the CD45/31-, EPCAM+, PDPN-negative, -low and -high cell populations depicted as a heatmap. Gene order is based on hierarchical clustering. C) Box plot showing the expression of genes associated with the ‘resting’ basal cell state and ‘differentiating’ basal cell state amongst CD45/31-, EPCAM+ PDPN-high (blue) and PDPN-low (green) cells. The box and whiskers represent the median and interquartile range. The points show biological replicates that fall outside of the interquartile range.

Gene expression differences between the PDPN-low and PDPN-high cell populations were less evident, but the highest expression of *TP63*, a gene associated with quiescent basal cells [8–10], was found in a subset of PDPN-high cells (Figure 4B). While two cell adhesion-related genes, *MPZL2* and *DST*, were identified within these samples (Figure 4B), no difference in cell adhesion to collagen VI was detected between PDPN-low and PDPN-high cells (Supplementary Figure 4).

Next, we assessed the expression of genes that have previously been found to distinguish quiescent, differentiating and cycling basal cells in single cell RNA sequencing studies. Gene lists were produced for each from a literature search (Supplementary Table 2). Upregulation of genes consistent with a differentiating basal cell profile was found in the PDPN-low cells, with higher levels of expression of *KRT19*, *NOTCH*-3, *KRT4*, *SERPINB3*, *HES1*, *SERPINB4*, *SCGB3A2*, *CEACAM5*, *CEACAM6* and *SERPINB13* than was found in PDPN-high cells (Figure 3C). In contrast, the PDPN-high cell population demonstrated higher expression of genes typically associated with the quiescent basal cell subset, including *TP63*, *NPPC*, *KRT15*, *DLK2* and *JAG2*. Genes associated with cycling basal cells were expressed at similar levels within the PDPN-low or PDPN-high cell populations (Supplementary Figure 5).

To validate these findings at the protein level, we compared TP63 expression between PDPN-low and PDPN-high cells. The two populations were sorted and cytospun onto slides for immunofluorescence analysis (Figure 5A). The PDPN-high population contained a significantly higher proportion of TP63-positive cells, 87.7%, than the PDPN-low population, 24.1% (Figure 5B). Moreover, when considering TP63-expressing cells only, cells in the PDPN-high group showed a significantly higher median fluorescence intensity of TP63 compared to cells in the PDPN-low group (Figure 5C). This finding supports transcriptomic analysis showing that basal cells in the PDPN-high group express higher levels of *TP63* than PDPN-low cells.

**Figure 5:**
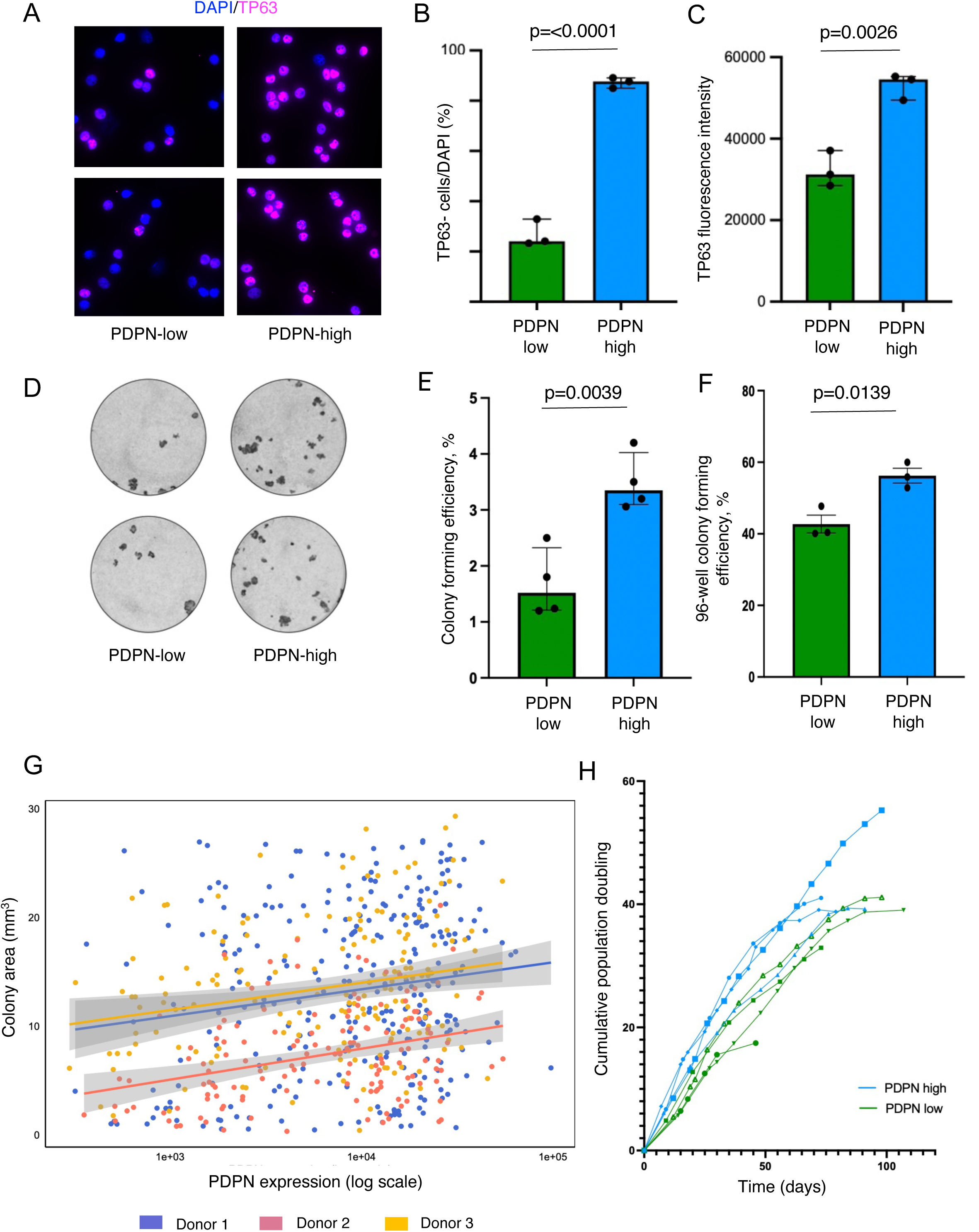
Higher PDPN expression amongst airway epithelial cells is associated with higher TP63 protein expression, higher colony-forming efficiency, larger colony size and shorter population doubling time. A) Immunofluorescence staining showing TP63 staining in DAPI+ nuclei from flow-sorted CD45/31-, EPCAM+, PDPN-low and PDPN-high cell populations. B) Bar chart showing the median proportion of TP63+/DAPI nuclei from flow-sorted CD45/31-, EPCAM+, PDPN-low and PDPN-high cell populations from three donors. The interquartile range is shown and individual points represent biological replicates. Unpaired t-test showed a significant difference, p<0.0001. C) Bar chart showing the median TP63 fluorescence intensity amongst CD45/31-, EPCAM+, PDPN-low and -high cells flow-sorted from three donors. The interquartile range is shown and individual points represent biological replicates. Unpaired t-test showed a significant difference, p=0.0026. D) Representative wells from a colony-forming assay showing two technical replicates from one proximal airway specimen. E) Median colony-forming efficiency (CFE) amongst CD45/31-, EPCAM+, PDPN-low and -high cells flow-sorted from four donors. The interquartile range is displayed and individual points represent biological replicates. Unpaired t-test showed a significant difference, p=0.0039. F) Mean CFE from single CD45/31-, EPCAM+, PDPN-low and –high cells that were individually plated. The level of PDPN expression was assessed retrospectively from index sorting of all seeded cells from three biological replicates. The standard error of the mean is shown. Unpaired t-test showed a significant difference, p=0.0139. (G) Scatterplot showing a significant positive correlation for each of the three biological replicates between PDPN expression of the seeding cell and area of the resulting colony (mm^2^). Colonies that did not grow (i.e. size = 0 mm^2^) were excluded from the analysis. The Pearson correlation co-efficient for each of the biological replicates is as follows; purple (r) = 0.116, p = 0.0491; pink (r) = 0.235, p = 0.0142 and orange (r) = 0.214, p = 0.0142. H) CD45/31-, EPCAM+, PDPN-high and –low cells were flow-sorted and cultured over serial passages to assess their population doubling time. The lines represent four biological replicates (blue = PDPN-high starting populations, green = PDPN-low populations).

### Higher PDPN expression is associated with basal stem cell potential *in vitro*

Finally, we investigated the progenitor cell potential of PDPN-low and PDPN-high basal cells *in vitro*. Colony-formation assays were performed to assess the clonogenic ability of individual basal cells. PDPN-low and PDPN-high cell populations were flow sorted and seeded at low density (1000 cells/well in a 6-well plate; Figure 5D). PDPN-high cells demonstrated a significantly higher colony-forming efficiency, 3.3%, than PDPN-low cells, 1.5% (Figure 5E). Similarly, when individual EPCAM+ epithelial cells were seeded into the wells of 96-well plates, PDPN-high cells had a significantly higher colony-forming efficiency, 56.2%, than PDPN-low cells, 42.7% (Figure 5F). In addition, colony size was weakly correlated with the level of PDPN expression on the seeding cell across three biological replicates (Figure 5G).

Freshly sorted PDPN-low and PDPN-high cells were both capable of long-term growth in cell culture and underwent serial passaging. PDPN-high cells showed a shorter population doubling time in comparison to PDPN-low cells (Figure 5H). However, Ki67 staining of freshly sorted PDPN-low and PDPN-high cells did not reveal a significant difference in cell proliferation after three days in the assay (Supplementary Figure 6A). To assess the differentiation potential of PDPN-low and PDPN-high cells, we performed tracheosphere cultures. In human bronchial epithelial cells (HBECs) that had been expanded in culture for one passage and fluorescence-activated cell-sorted into PDPN-low and PDPN-high populations, the tracheosphere-forming capacity was comparable between PDPN-low and PDPN-high cells (Supplementary Figure 7B). Both populations formed tracheospheres, and these contained KRT5+ basal cells, MUC5AC+ mucosecretory cells and acetylated tubulin (ACT+) ciliated cells (Supplementary Figure 6C).

## Discussion

Our study shows that PDPN can be used to mark KRT5+ basal epithelial cells with a balance of sensitivity and specificity that is superior to that of ITGA6 or NGFR. PDPN expression levels also allowed us to distinguish transcriptomically and functionally distinct cell populations. While both PDPN-low and PDPN-high cell populations expressed KRT5, PDPN-low cells expressed genes associated with differentiating basal cells, such as *KRT4, KRT19* and *NOTCH3*, while PDPN-high cells expressed high levels of genes associated with basal cell quiescence, such as *TP63*. We confirmed that TP63 was more highly expressed in PDPN-high cells at the protein level, and functional data were consistent with this interpretation: PDPN-high cells exhibited a higher colony-forming efficiency and generated larger colonies than PDPN-low cells *in vitro*.

Our findings in human airway basal cells are reminiscent of the heterogenous potential of epidermal keratinocytes. When single basal keratinocytes are plated to form colonies, resultant clones can be classified, according to their appearance, as either holoclones, meroclones or paraclones [28]. Holoclones are keratinocytes capable of extensive self-renewal and proliferation – the colonies produced are considered to derive from stem cells. Meroclones have a limited proliferative capacity and paraclones contain keratinocytes with a very limited lifespan, which gives rise to small, terminal colonies; these keratinocytes are thought to represent early and late-stage transit-amplifying cells, respectively [29]. TP63 is highly expressed in holoclones, but absent from paraclones [30]. Overall, our data support a model in which PDPN-high basal cells represent the basal stem cell population with the greatest regenerative potential—akin to the ‘quiescent’ or ‘resting’ basal cells identified in airway single cell RNA sequencing studies and Green’s holoclones [8, 11–12]. Meanwhile, PDPN-low basal cells may represent a more differentiated basal cell population—similar to the ‘differentiating’ or ‘suprabasal’ cells observed in transcriptomic datasets, and equivalent to Green’s meroclones and/or paraclones [11–13].

PDPN is a ligand for C-type lectin-like receptor 2 (CLEC2) and can exert effects on cellular signaling [31]. PDPN can also act as a receptor, for example, for galectin 8 [32] or CCL21 [33]. Pathologically, PDPN is upregulated in some human cancers, including lung squamous cell carcinoma [34] and mesothelioma [35]. *In vitro*, PDPN expressed on lung squamous cell cancer cells binds CLEC-2 on platelets to promote their aggregation, which in turn enhances tumor cell growth [36]. In epithelial cells, the consequences of PDPN loss seem to be context dependent; no defects in wound healing were observed following epithelial-specific deletion in mouse skin [37], but significant defects in mouse mammary stem cell function were seen following PDPN deletion [38]. PDPN plays a role in lung development as *Pdpn*^-/-^mice have impaired alveolus formation and die from respiratory failure shortly after birth [39]. However, it is unclear whether PDPN plays a functional role in airway basal cells. Our data justify mechanistic investigation of PDPN’s role in basal cell stemness and airway epithelial cell differentiation. It is also plausible that PDPN may influence cell adhesion and migration based on known roles for the protein in other tissue types.

Our results support the use of PDPN as a cell surface marker to identify airway basal stem cells, with the expression of PDPN decreasing during airway epithelial differentiation. The ability to select airway basal cells, whilst maintaining their viability for functional analysis and downstream use will allow in-depth study of basal cell behavior *in vitro*, as well as opening the door for future clinical applications. Selecting airway basal cells with the highest *in vitro* regenerative potential may have potential applications in regenerative medicine, for example in airway cell therapies requiring *in vitro* expansion [40, 41].

### Conclusion

Our results demonstrate the functional heterogeneity of human airway basal cells in *ex vivo* cultures. PDPN is an effective surface marker of basal cells and the level of its expression demarcates subpopulations with transcriptomic and functional differences. We propose that PDPN-high cells have a higher level of stemness and growth potential than PDPN-low cells and that these differences reflect the airway differentiation process.

## Materials and methods

### Human proximal airway samples

Patients undergoing lobectomy, primarily for lung cancer treatment, were recruited and consented under the ethics ‘An Investigation into the Molecular Pathogenesis of Lung Disease II’ (IRAS: 245471, REC ref: 18/SC/0514). Second-to fourth-generation macroscopically normal airways, situated away from the tumor site, were collected. Patient characteristics are summarized in Supplementary Table 1.

### Single cell RNA sequencing analysis

This scRNAseq dataset comprises a total of 41,730 epithelial cells from the trachea. Data from two publicly accessible datasets (GSE134174, *n* = 12 donors & EGAS00001004082, *n* = 9 donors) [9, 51] were combined with data from six tracheal biopsies that were processed and sequenced in-house (GSE276610, *n*= 6 donors) [16]. Data visualization was performed using Scanpy (version 1.10.1), where UMAP projections and dot plots for selected markers (*PDPN*, *KRT5* and *TP63*) were generated to visualize cellular heterogeneity and assess marker expression across different epithelial cell populations.

### Histology

Proximal airway samples were fixed overnight in 4% paraformaldehyde (PFA) and dehydrated through an ethanol gradient using a Leica TP 1050 vacuum tissue processor. The tissues were embedded in paraffin and sectioned at 4 µm thickness with a microtome. Hematoxylin and eosin (H&E) staining was performed on an automated system (Tissue-Tek DRS, Sakura). Stained slides were scanned with a Nanozoomer Whole Slide Imager (Hamamatsu Photonics) and analyzed using NDP.View2 software.

### Immunofluorescence staining

FFPE slides were de-waxed and underwent heat induced epitope retrieval (10 mM sodium citrate buffer, pH 6.0) at 95°C for 15 minutes. After cooling, slides were washed with phosphate buffered saline (PBS). Slides were incubated in 3% H_2_O_2_ for 10 minutes at room temperature (RT), washed twice with PBS, then blocked for one hour in a PBS solution containing 5% donkey serum, 3% BSA and 0.25% Triton X-100. Primary antibodies were diluted in the blocking buffer, then incubated overnight at 4°C (chicken anti-KRT5, 1:500, BioLegend, 905904; rat anti-PDPN IgG2a, 1:50, MBL D189-1). Slides were washed twice with phosphate buffered saline with Tween (PBST). For the PDPN staining, amplification was required using the TSA signal amplification kit (PerkinElmer). Slides were incubated in Rat-HRP antibody (A10549, Invitrogen, 1:250) for 45 minutes at RT. Slides were then washed twice in PBST (0.1% Triton PBS) and incubated with biotin diluted in amplification buffer for eight minutes. Following 2 washes with PBST, slides were incubated in Streptavidin HRP antibody (S32355, Invitrogen, 1:250) for 30 minutes at RT. Slides were incubated with secondary antibody, AF 488 conjugated polyclonal goat anti-chicken antibody (A32931, Invitrogen, 1:500) diluted in block solution either at RT for three hours or overnight at 4°C. After antibody staining, nuclei were stained with DAPI (1 μg/ml stock, Invitrogen, 1:1000 in PBS) and sections were mounted with Fluoromount G mounting medium (Invitrogen). Images were acquired using a Leica DMi8 or a Zeiss LSM880 confocal microscope and processed with Fiji software.

### Enzymatic digestion of airway tissue

Airway samples were manually homogenized and digested sequentially. First, tissue was digested with 16 U dispase (Corning) in RPMI media (Gibco) at RT for 20 minutes. The digested tissue was then transferred to Eppendorf tubes containing 0.1% trypsin (Sigma) and incubated at 37°C for 30 minutes. The remaining dispase digestion was quenched with 200 μl fetal bovine serum (FBS) and placed at 4°C. After 30 minutes, Eppendorf tubes from both the first and second digestion stages were combined. In some cases, liberase TL (Roche) replaced dispase and trypsin. Tissue was incubated with 150 μg/ml liberase TL in RPMI medium at 37°C for 30 minutes with gentle agitation. Digested material was filtered through a 100 μm cell strainer (Corning) to obtain single-cell suspensions.

### Fluorescent-activated cell sorting (FACS)

Single-cell suspensions of fresh airway tissue or cultured HBECs were pelleted and resuspended in sorting buffer (PBS, 1% FBS, 25 μM HEPES, 1 mM EDTA). Samples were incubated with the following antibodies (all 1:50) for 15 minutes at 4°C: CD31 (BV421; BioLegend; 303124), CD45 (BV421; BioLegend; 304031), EPCAM (APC; BioLegend; 324208) and/or PDPN (PE-Cy7; BioLegend; 337013). Controls included unstained cells, single stained cells or beads (Invitrogen), and fluorescence-minus-one (FMO) controls. 2 μl DAPI (1 μg/ml stock, Invitrogen; 1:100 in PBS) was added as required. Samples were sorted using a BD FACSAria Fusion sorter at the UCL Cancer Institute Flow Cytometry Core Facility. Sorted cells were collected into epithelial cell culture medium with Y-27632.

### Flow cytometric analysis

Single cell suspensions of fresh airway cells were stained with the antibodies above, plus those against ITGA6 (PerCP/Cy5.5; BioLegend; 313617) and NGFR (PE; BioLegend; 345105) both at 1:200. For KRT5 flow cytometry experiments, cells were fixed in CellFix buffer (BD BioSciences) for ten minutes and permeabilized with saponin permeating solution (Thermo Fisher Scientific). Anti-KRT5 (conjugated to Alexa Fluor 488; Abcam; 193894) was then added at 1:200, followed by DAPI, as above. Cells were evaluated using a BD LSR Fortessa flow cytometer and analyzed using FlowJo (Tree Star Inc, USA).

### Bulk RNA sequencing of flow sorted cell populations

Freshly sorted cells from three populations (CD45/31-, EPCAM+, PDPN-negative, -low or -high) were collected into Eppendorf tubes containing epithelial cell culture medium with Y-27632 and resuspended in cell suspension buffer (0.9 ml of 1x PBS/10% BSA; 0.1 ml of 0.5 M EDTA) once transported to the laboratory. RNA was extracted and isolated using the Arcturus PicoPure RNA Isolation Kit (Life Technologies Ltd) according to the manufacturer’s instructions. Quantity and quality of samples was assessed using a NanoDrop spectrophotometer and Qubit high sensitivity kit (Thermo Fisher Scientific). The cells were sequenced at the UCL Genomics Core Facility; libraries were created using the SMARTer Stranded Total RNAseq Kit (Clontech) and quality control analysis of RNA integrity (QC) was performed with High Sensitivity RNA ScreenTape using TapeStation Analysis software (Agilent Technologies). Illumina RNAseq was performed using 0.5x NextSeq for 75 cycles (Illumina; 43PE, ∼33M reads per sample).

### Bulk RNA sequencing analysis

Following sequencing, run data were demultiplexed and converted to fastq files using Illumina’s bcl2fastq Conversion Software v2.19. Quality control and adapter trimming were performed using fastp [45] version 0.20.1 with default settings. Fastq files were then tagged with the UMI read (UMITools [46]) and aligned to the human genome UCSC hg38 using RNA-STAR [47] version 2.5.2b. Aligned reads were UMI deduplicated using Je-suite [48] version 1.2.1 and count matrices were obtained using featureCounts. Downstream analysis was performed using the R statistical environment version 3.5.0 with Bioconductor version 3.8.0 [48,49]. Counts were compared between the three populations with DESeq2 [48] using the default settings. Heatmaps were plotted using the pheatmap package, implementing a complete linkage clustering method. All other plots were created using ggplot2. The gene lists containing markers for resting, differentiating and terminal basal cells (Supplementary Figure 2) were identified from review of the literature.

### Human airway epithelial cell culture

3T3-J2 mouse embryonic fibroblasts (a kind gift from Prof Fiona Watt, King’s College London, U.K.) were cultured and expanded in DMEM plus 9% bovine calf serum and 1x penicillin–streptomycin, as previously described [42]. 3T3-J2 cells were inactivated by 0.4 μg/ml mitomycin C treatment (Sigma-Aldrich) and seeded at a density of 20,000 cells/cm^2^ a day prior to the addition of epithelial cells. Epithelial cell culture medium with Y-27632 consisted of DMEM/F12 in a 3:1 ratio containing 1x penicillin–streptomycin (Gibco), 5% fetal bovine serum (Gibco) supplemented with 5 μM Y-27632 (Cambridge Bioscience), hydrocortisone (25 ng/ml; Sigma-Aldrich), epidermal growth factor (0.125 ng/ml; Sino Biological), insulin (5 μg/ml; Sigma-Aldrich), 0.1 nM cholera toxin (Sigma-Aldrich), amphotericin B (250 ng/ml; Thermo Fisher Scientific), and gentamicin (10 μg/ml; Gibco), as previously described [41].

### TP63 expression in cytospun cell populations

Freshly flow-sorted CD45/31-, EPCAM+, PDPN-low and PDPN-high cells were resuspended in 200 μl PBS and centrifuged at 800 rpm for five minutes onto poly-L-lysine-coated adhesive slides. Slides were air-dried for ten minutes and fixed in 2% PFA at RT for 15 minutes, washed in PBS and a hydrophobic ring was drawn around the sample using an ImmEdge pen (Vector Laboratories). Sections were blocked with 1% bovine serum albumin (Merck), 5% normal goat serum (Abcam) and 0.1% Triton X-100 (Sigma-Aldrich) in PBS for one hour at RT. An anti-TP63 antibody (ab124762, Abcam; 1:400) was diluted in block buffer and applied to slides overnight at 4°C. Secondary antibody conjugated to species appropriate Alexa Fluor dyes was diluted 1:1,000 in block buffer and applied to slides for two hours at RT in the dark. DAPI (1 μg/ml stock, Invitrogen; 1:10,000 in PBS) was applied to the slides for 20 minutes. Slides were washed twice in PBS and a coverslip was applied manually with Immu-Mount (Thermo Fisher Scientific). Images were acquired using a Leica DMi8 fluorescence microscope.

### Colony formation assays

Culture plates were rat tail collagen I-coated (Corning; 50 µg/ml in sterile 0.02N acetic acid for two hours at RT), washed with PBS and 3T3-J2 feeder cells were added as above one day in advance. For the 6-well plate assay, 1000 freshly sorted CD45/31-, EPCAM+, PDPN-low or PDPN-high cells were plated in triplicate. After 10 days, feeder cells were removed by differential trypsinization, the colonies were fixed and then stained with crystal violet solution (Sigma Aldrich). Wells were imaged using a Zeiss Airyscan 880 microscope and colony forming efficiency calculated. For the 96-well plate assay, fresh CD45/31-, EPCAM+ cells were individually sorted into wells and their level of PDPN expression was recorded by index sorting. After 14 days, wells containing colonies of >10 cells were counted and imaged using a Leica DMi8 microscope and colony area was calculated using Fiji.

### Ki67 staining

8-well chamber slides (Ibidi) were prepared with collagen I coating and inactivated 3T3-J2 cells as outlined above. 8000 freshly sorted CD45/31-, EPCAM+ PDPN-low or PDPN-high cells were plated in triplicate in epithelial cell culture medium with Y-27632 onto the chamber slide wells. After three days, wells were washed with sterile PBS and fixed with 4% PFA. Cells were incubated with rabbit anti-Ki67 antibody at 1:500 (Thermo Fisher Scientific) overnight at 4°C, prior to washing and incubating with goat AF488-conjugated anti-rabbit IgG at 1:500 (Thermo Fisher Scientific) for three hours. DAPI (1 μg/ml stock, Invitrogen; 1:10,000 in PBS) was added and staining was quantified using a Zeiss Airyscan 880 microscope and ImageJ software. Four images were taken of each of the replicate wells and the number of Ki67+ cells / number of DAPI+ cells * 100 was calculated and averaged.

### Population doublings

Comparative expansion rates of CD45/31-, EPCAM+ PDPN-low or PDPN-high basal cells were evaluated by population doubling time. Identical numbers of sorted cells were seeded and parallel cultures were expanded over serial passages in T25 flasks, with passaging once 80% confluence was reached. Immediately prior to passage, cell counts using a manual hemocytometer were recorded and 125,000 cells were re-seeded in T25 flasks containing fresh 3T3-J2 feeder layers. Cultures were maintained for 100 days, or until cultures failed. Population doublings (PD) were calculated as PD = 3.32 * (log (cells harvested / cells seeded), 10) and were plotted against time in days. Doubling times were calculated by interpolating from the line of best fit and subtracting time (PD20) from time (PD10).

### Tracheosphere assay

96-well ultra-low attachment plates (Corning) were coated with 30 μl tracheosphere medium containing 25% growth factor-reduced Matrigel (BD Sciences). Tracheosphere medium was as previously described [42]. Primary cultured cells were flow sorted into CD45/31-, EPCAM+ PDPN-low or PDPN-high populations. 2500 cells from each population were resuspended in 65 μl tracheosphere medium with 5% Matrigel and 5 μM Y-27632 and plated with six replicates. On days 3, 8 & 14, each well was fed by addition of a further 70 μl tracheosphere medium. After 21 days, tracheospheres were fixed; half of the replicates were processed for whole mount staining, and the other half paraffin embedded. For whole mount staining, tracheospheres were resuspended in a blocking solution (PBS with 0.1% Triton X100, 1% DMSO, 1% BSA and 1% goat serum). The following primary antibodies were added; MUC5AC (mouse IgG1, Sigma) 1:200, KRT5 (chicken, BioLegend) 1:500, and acetylated tubulin (ACT; mouse IgG2b, Sigma) 1:500, and the samples were incubated for 40 hours with gentle agitation at 4°C. The tracheospheres were then washed and incubated with secondary antibodies; anti-rabbit IgG AF488 (Thermo Fisher Scientific) 1:500, anti-mouse IgG2b AF647 (Thermo Fisher Scientific) 1:500 and anti-mouse IgG1 AF555 (Thermo Fisher Scientific) 1:500 diluted in PBS with 0.1% BSA for three hours. The tracheospheres were then washed and DAPI (1 μg/ml stock, Invitrogen; 1:10,000 in PBS) added. Following this they were mounted on a glass slide with separators using Vectashield (Vector). Immunofluorescent whole-mount images were taken on the Zeiss Airyscan 880 and analyzed using Fiji. For paraffin embedding, the tracheospheres were resuspended in PBS followed by HistoGel at 65°C (Thermo Fisher Scientific) which was allowed to cool and solidify. The gel discs were dehydrated through an ethanol gradient prior to being paraffin embedded. The paraffin embedded blocks were cut and stained using the antibodies outlined above.

### Adhesion assay

96-well plates were coated with collagen IV from human placenta (Sigma-Aldrich) at 250 µg/ml [44]. 2000 cells freshly sorted CD45/31-, EPCAM+ PDPN-low or PDPN-low populations were plated in triplicate and incubated for 30 minutes at 37°C. After 30 minutes, unbound cells were removed by washing wells three times with PBS. Wells were fixed with 4% PFA and stained with DAPI. Wells were imaged using the Leica microscope and DAPI+ cells were manually counted.

### Statistics

Statistical analysis was performed using GraphPad Prism 10. Variables are frequently presented as median values, of three technical replicates, with the interquartile range displayed. Biological replicates represent airway samples taken from different donors. Group differences were compared using an unpaired t-test and where more than two groups were being compared, an ANOVA test was run in the first instance.

## Acknowledgements

The authors thank Dr Sophie Acton (University College London) for sharing her knowledge and expertise in podoplanin biology. We acknowledge the roles of Mr David Lawrence, Mr Nikolaos Panagiotopoulos, Mr Sofoklis Mitsos and Mr Achilles Antonopoulos (Cardiothoracic Surgery, University College Hospital, Westmoreland Street), along with the patients who took time to read and hear about our study. We thank Dr Ricky Thakrar & Bernadette Carroll for their additional help with research airway tissue sampling, Dr David Moore, Dr Elaine Borg, and Dr Mary Falzon for their assistance with dissection of lobectomy specimens and histology (Pathology Department, University College Hospital London), and Yanping Gui and George Morrow (UCL Cancer Institute Flow Cytometry Core Facility, University College London) and Jamie Evans (Division of Medicine, University College London Flow Cytometry Core Facility, Rayne Building, University College London) for assistance with flow cytometry and FACS experiments. Finally, we thank Tony Brooks and Dr Paola Niola (UCL Genomics, University College London) for performing bulk RNA sequencing, and Simon Broad and Professor Fiona Watt (King’s College London, UK) for providing 3T3-J2 fibroblasts.

## Funding

This work was funded by NIHR (award reference NIHR-INF-0383) and supported by grants from the Longfonds BREATH lung regeneration consortium (to S.M.J.) and the UK Regenerative Medicine Platform (UKRMP2) Engineered Cell Environment Hub [Medical Research Council (MRC); MR/R015635/1] (to R.E.H. and S.M.J.). R.E.H. was supported by a Wellcome Trust Sir Henry Wellcome Fellowship (WT209199/Z/17/Z), a NIHR Great Ormond Street Hospital BRC Catalyst Fellowship, GOSH Charity (V4322), and the Royal Society (RG\R1\241421). S.M.J. and R.E.H receive support from the CRUK Lung Cancer Centre of Excellence (C11496/A30025). S.M.J. is supported by a CRUK programme grant (EDDCPGM\100002) and an MRC programme grant (MR/W025051/1), the CRUK City of London Centre, the Rosetrees Trust, the Roy Castle Lung Cancer foundation, the Garfield Weston Trust, and University College London Hospitals Charitable Foundation. S.M.J.’s work is supported by a Stand Up To Cancer-LUNGevity Foundation American Lung Association Lung Cancer Interception Dream Team Translational Research Grant and Johnson and Johnson (SU2C-AACR-DT23-17). Stand Up To Cancer is a division of the Entertainment Industry Foundation. Research grants are administered by the American Association for Cancer Research, the Scientific Partner of SU2C. M.J.R. was supported by a CRUK City of London Centre Clinical Academic Training Fellowship (BCCG1C8R). This work was partly undertaken at UCL/UCLH and partly at UCL ICH/GOSH who received a proportion of funding from the Department of Health’s NIHR Biomedical Research Centre’s funding scheme.

## Author Contributions

Conception and design: S.E.C., R.E.H., K.H.C.G. and S.M.J.; Provision of study materials or patients: S.E.C., C.C.P., S.G-L., M.J.R., M-B.E-M., Z.C.H., K.E.J.O., C.R.K., H.H., A.A.B. and K.H.C.G.; Investigation: S.E.C., C.C.P., Z.E.W., Y.I, S.G-L., M.J.R., J.C.O., M-B.E-M., R.E.H. and K.H.C.G. Data analysis and interpretation: S.E.C., A.P., Y.I., H.S-C., M-B.E-M., R.E.H. and K.H.C.G. Manuscript writing: S.E.C., A.P., Z.E.W., Y.I., M.J.R.,. E.H. and K.H.C.G. Manuscript review & editing: Y.I., S.G.-L., R.E.H., and S.M.J. Funding acquisition: S.E.C., R.E.H., K.H.C.G. and S.M.J. Supervision: Y.I., S.G-L., M-B.E-M., R.E.H., K.H.C.G and S.M.J.

## Competing Interests

S.M.J. has received fees for advisory board membership from BARD1 Life Sciences. S.M.J. has received grant income from GRAIL Inc. and is an unpaid member of a GRAIL advisory board. S.M.J. has received lecture fees for academic meetings from Chiesi and AstraZeneca. R.E.H. has received lecture fees for academic meetings and consultancy fees from AstraZeneca. None of the other authors has any conflicts of interest, financial or otherwise, to disclose.

## Supplementary Tables

**Supplementary Table 1:** Patient characteristics of the airway samples used within the study.

| ID | Age (year) | Sex | Procedure | Indication for procedure | Sites of sampling | Experiment | Medical history of lung disease | Smoking Status | Pack-year | Duration of smoking cessation (year) |
| --- | --- | --- | --- | --- | --- | --- | --- | --- | --- | --- |
| P313 | 53 | F | Right upper completion lobectomy | PET avid lesion | 2nd gen airway from right upper lobe | CFA | Nil | Current smoker | 40 | N/A |
| P326 | 59 | F | Right pneumonectomy | Lung adenocarcinoma | 2nd gen airway | CFA | Nil | Ex-smoker | 40 | <1 |
| P332 | 74 | F | Right upper completion lobectomy | PET avid lesion | 2nd gen airway | RNAseq | Nil | Current smoker | 45 | N/A |
| P334 | 77 | M | Left upper lobectomy | Squamous cell lung carcinoma | 2nd gen airway | RNAseq & CFA | Nil | Ex-smoker | 50 | 10 |
| P339 | 78 | F | Left upper lobectomy | Lung adenocarcinoma | 2nd gen airway | RNAseq | COPD | Ex-smoker | 40 | 10 |
| P341 | 75 | M | Left upper lobectomy | Squamous cell lung carcinoma | 2nd gen airway | RNAseq | Nil | Ex-smoker | 10 | 50 |
| P345 | 74 | M | Right lower lobectomy | Squamous cell lung carcinoma | 2nd gen airway | RNAseq | Nil | Ex-smoker | 10 | 20 |
| P350 | 83 | M | Bronchoscopic biopsy | CIS surveillance | Main carina | RNAseq | Previous left lower lobectomy for lung cancer | Ex-smoker | 45 | 40 |
| P360 | 72 | M | Right upper lobectomy | Lung adenocarcinoma | 2nd gen airway | RNAseq & CFA | Nil | Never smoker | N/A | N/A |
| P362 | 83 | M | Right lower lobectomy | PET avid lesion | 2nd gen airway | RNAseq | Nil | Ex-smoker | 45 | 10 |
| P363 | 76 | M | Right upper & middle lobectomies | Lung adenocarcinoma | 2nd gen airway | CFA | Nil | Ex-smoker | 38 | <1 |
| P364 | 83 | F | Left lower lobectomy | Lung adenocarcinoma | 2nd gen airway | RNAseq | Nil | Never smoker | N/A | N/A |
| P366 | 68 | M | Right upper lobectomy | Lung adenocarcinoma | 2nd gen airway | Basal cell marker flow analysis | Nil | Never smoker | N/A | N/A |
| P372 | 58 | F | Left upper lobectomy | Squamous cell lung carcinoma | 2nd gen airway | Tracheosph ere assay | COPD | Current smoker | 44 | N/A |
| P376 | 69 | M | Right upper lobectomy | PET avid lesion | 2nd gen airway | Tracheosph ere assay & CFA | Asthma | Ex-smoker | 25 | 5 |
| P377 | 70 | M | Right upper lobectomy | Squamous cell lung carcinoma | 2nd gen airway | Tracheosph ere assay & population doubling | COPD | Ex-smoker | 56 | <1 |
| P378 | 55 | F | Completion lobectomy | PET avid lesion | 2nd gen airway | CFA & population doubling | Nil | Ex-smoker | 30 | 1 |
| VM | 49 | M | Bronchoscopic biopsy | Surveillance for lung papilloma | Carina | Basal cell marker flow analysis | Papilloma of LMB, mild emphysema, Langerhans | Current smoker | 10 | N/A |
| MF | 56 | M | Therapeutic ML | VC granuloma | Trachea | Basal cell marker flow analysis | Asthma | Never smoker | N/A | N/A |
| LH | 68 | F | Therapeutic ML | VC polyp | Trachea | Basal cell marker flow analysis | Nil | Never smoker | N/A | N/A |

**Supplementary Table 2:** Categorization of literature single cell RNA sequencing basal cell clusters. The genes that were expressed within these have been used to interrogate the flow-sorted PDPN populations in Figure 2.

| scRNAseq Basal Cell Cluster Terminology | Broad Basal Cell Category | Differential genes expressed | scRNAseq Literature Reference |
| --- | --- | --- | --- |
| 'Non-cycling basal cells'<br>'Basal-1 cells'<br>'Basal resting cells' | Resting | *TP63, NPPC, KRT15, KRT17, DLK2* | Plasschaert et al., Ruiz-Garcia et al., Vieira-Braga et al., Sikkema et al., Deprez et al. |
| 'Basal luminal progenitor cells'<br>'Suprabasal cells'<br>'Basal 2 cells'<br>'Differentiating basal cells' | Transitional | *KRT19, KRT4,, SERPINB3, HES1, SERPINB4, SCGB3A2* | Plasschaert et al., Ruiz-Garcia et al. |
| 'Cycling basal cells'<br>'Hillock-like; highly proliferative cells'<br>'Proliferating basal cells' | Cycling | *MKI67, KRT8, KRT14, KRT13, TOP2A, DSG3, RAB38, TN54* | Plasschaert et al., Ruiz-Garcia et al., Vieira-Braga et al., Sikkema et al., Travaglini et al. |

## Supplementary Figures

**Supplementary Figure 1:**
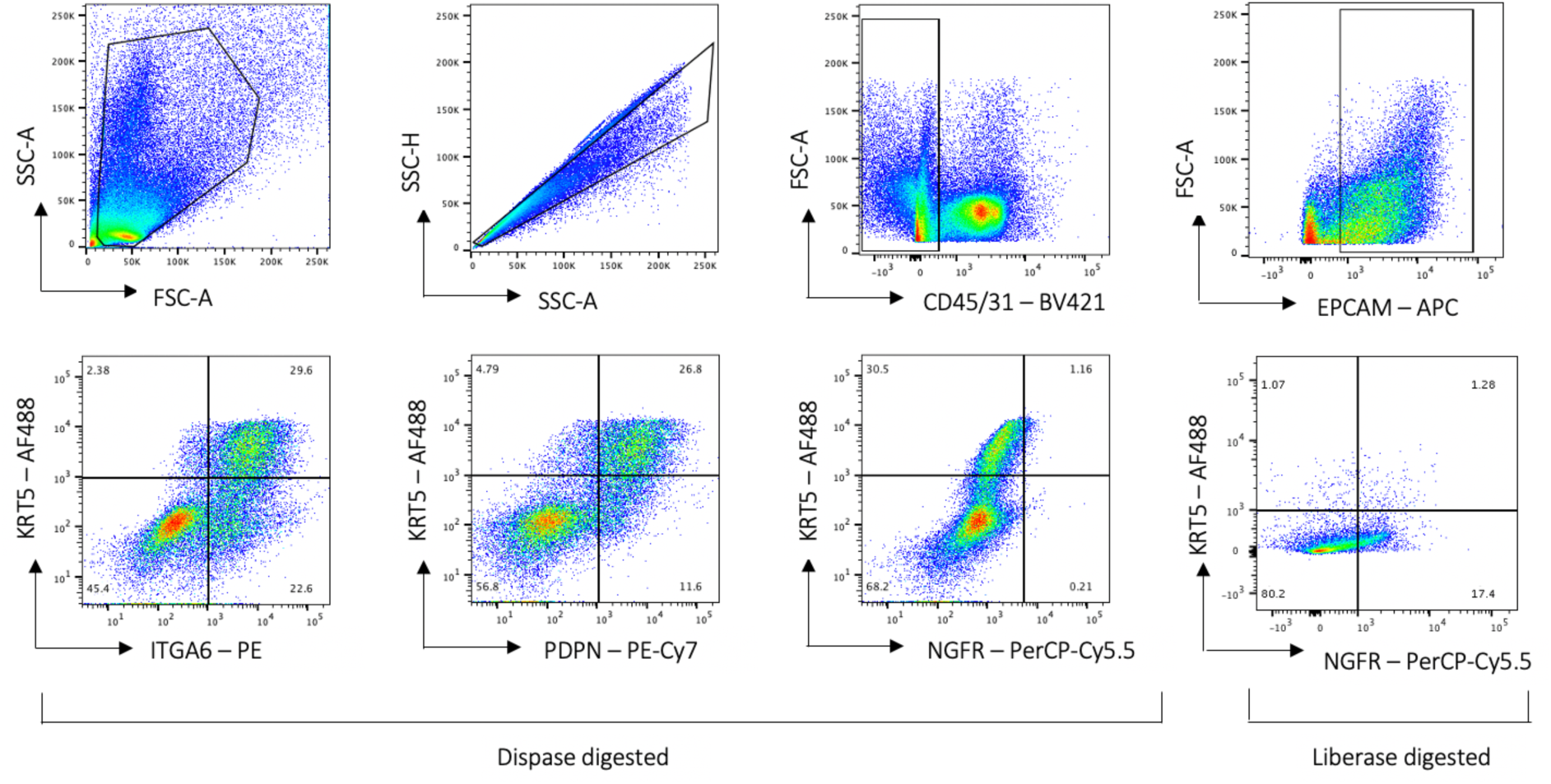
Flow sorting strategy for airway basal cells. Representative flow cytometric analysis demonstrating the expression of candidate extracellular basal cell markers (ITGA6, PDPN and NGFR) among CD45/31-, EPCAM+, KRT5+ and KRT5-proximal airway cells. Cells were digested with dispase and trypsin. In view of the low sensitivity result obtained for NGFR, thought to be a result of dispase cleavage, liberase digestion was performed on a further two replicates for which an isolated flow cytometry plot is shown - the initial sorting strategy was identical.

**Supplementary Figure 2:**
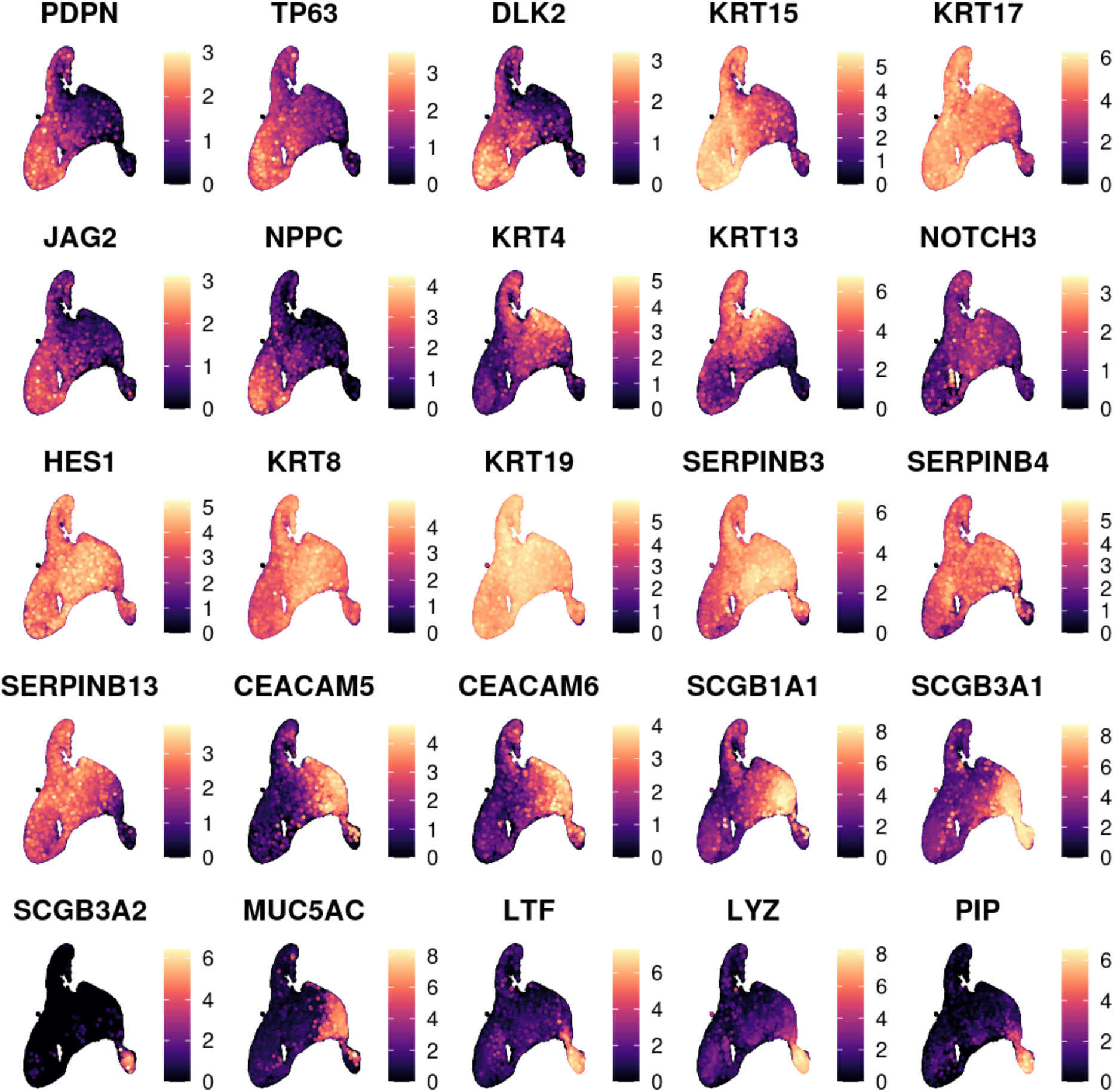
Extended visualization of selected gene expression in RNA sequencing dataset. UMAP visualization of gene expression of resting, differentiating and terminal markers.

**Supplementary Figure 3:**
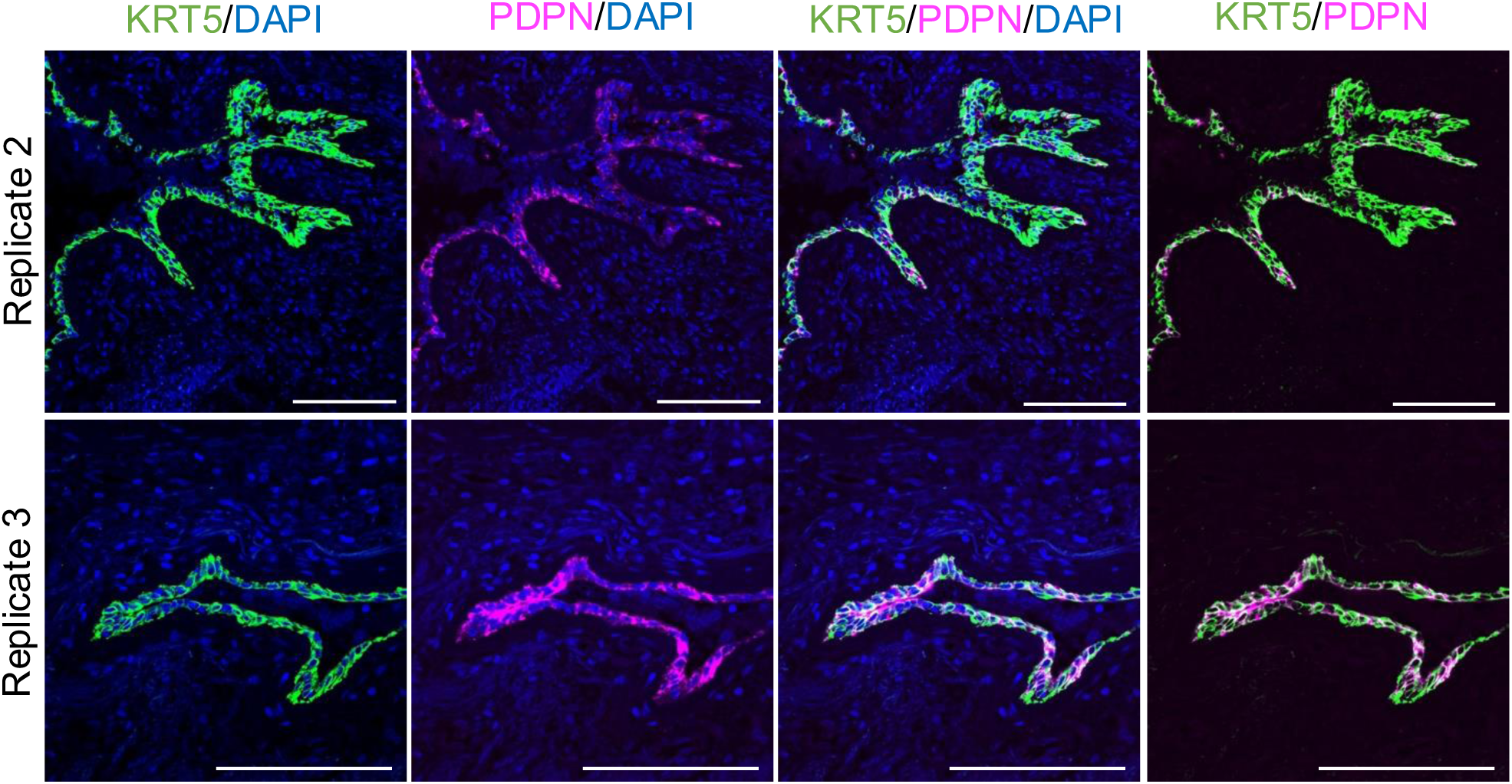
Extended immunofluorescence data showing KRT5 and PDPN expression in human airway samples. Related to Figure 2F; two additional donor samples are shown.

**Supplementary Figure 4:**
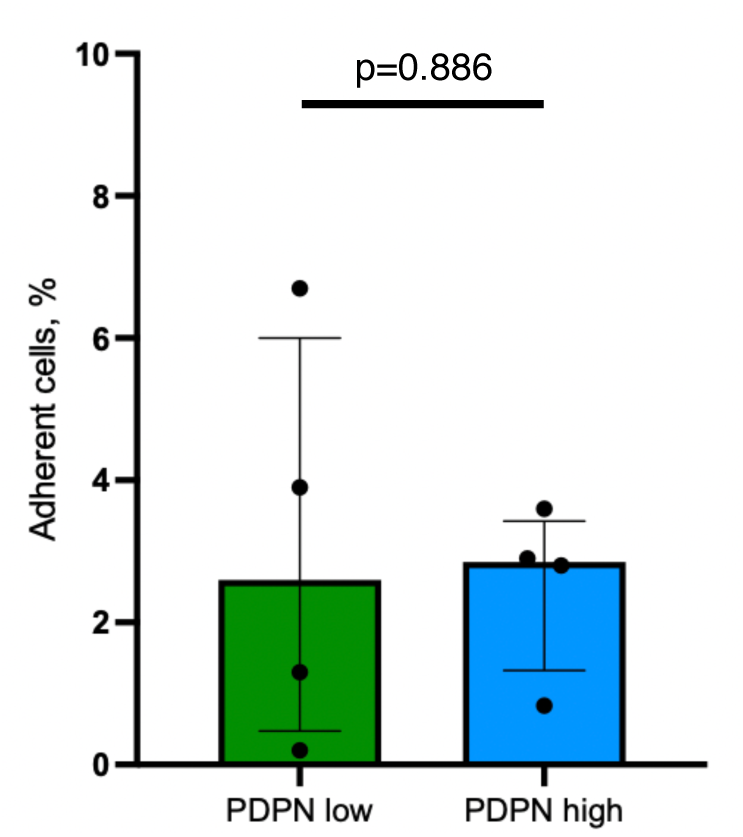
Cell attachment assay comparing PDPN low and PDPN high cells sorted from cultured airway basal cells. Bar chart showing the median percentage of adherent CD45/31-, EPCAM+, PDPN-low and -high cells that remained attached to the plate after 30 minutes of incubation followed by treatment with sequential washes. The error bars indicate the interquartile range. Each point represents a biological replicate. An unpaired t-test was performed for which the p-value is shown.

**Supplementary Figure 5:**
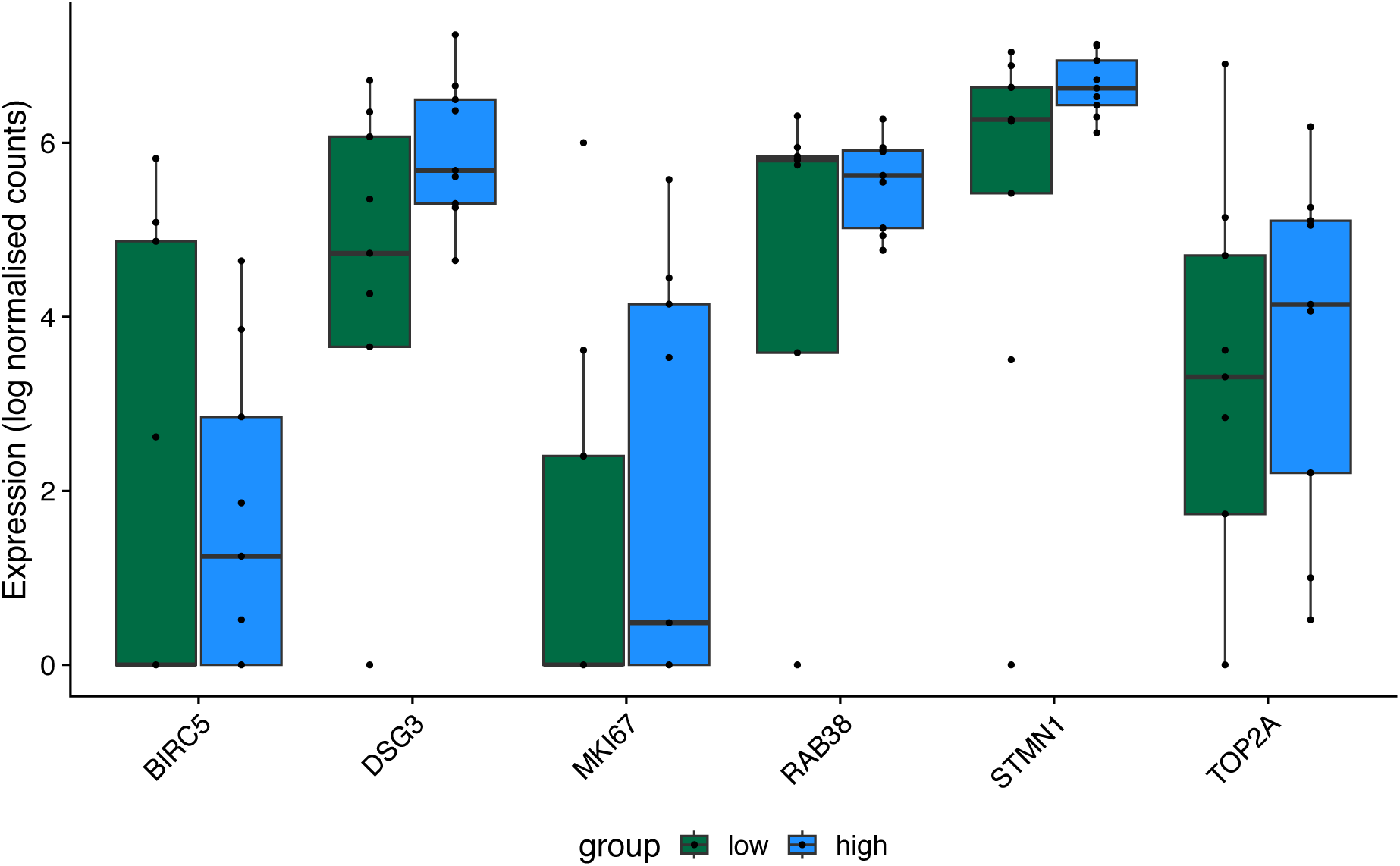
‘Cycling’ basal cell gene expression analysis. Box plot showing the expression of genes associated with the ‘cycling’ basal cell state amongst CD45/31-, EPCAM+ PDPN-high (blue) and PDPN-low (green) cells. Relevant genes were identified from a literature search. The box and whiskers represent the median and interquartile range. Each point represents a biological replicate.

**Supplementary Figure 6:**
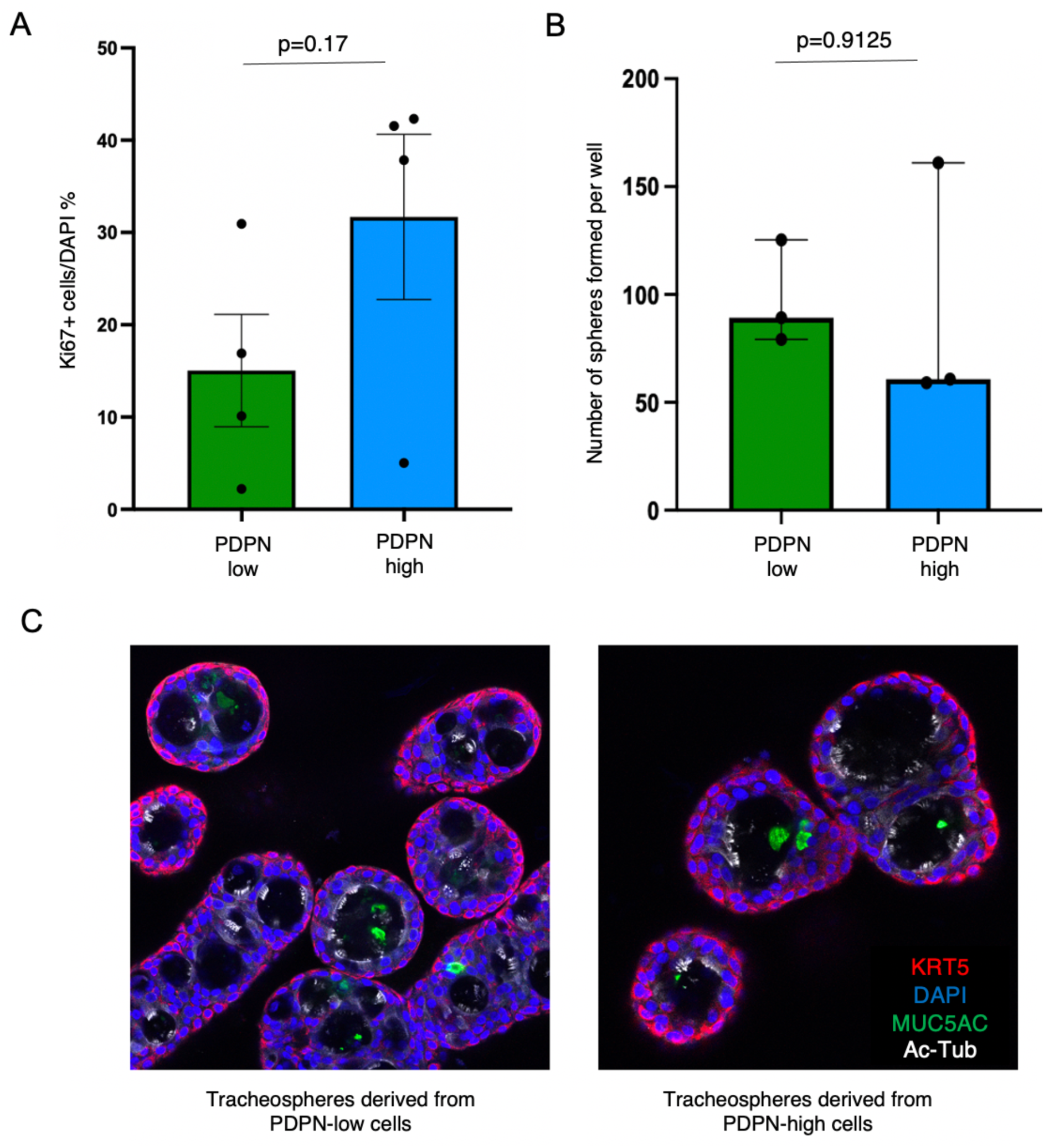
Extended functional analysis comparing the behavior of PDPN low and PDPN high cells in *in vitro* assays. A) Bar chart comparing the mean Ki67+ cells, % between PDPN-high and –low cells. Error bars indicate the standard error of the mean. Each point represents an individual biological replicate. An unpaired t-test was non-significant, p=0.17. B) Bar chart showing the median number of spheres that were formed per well for CD45/31-, EPCAM+, PDPN-low and –high cells. The bar represents the interquartile range and each point represents a biological replicate. An unpaired t-test was non-significant, p=0.9125. C) Immunofluorescence staining images of the tracheospheres after 21 days in culture from PDPN-low and PDPN-high cell populations. Tracheospheres were fixed and whole-mount stained with antibodies to KRT5 (red), MUC5AC (green), acetylated tubulin (ACT; white) & DAPI (blue).

